# Spindle alignment is uncoupled with Kar9 symmetry breaking

**DOI:** 10.1101/2020.09.01.278523

**Authors:** Miram Meziane, Rachel Genthial, Jackie Vogel

**Affiliations:** Department of Biology, McGill University, Montreal QC H3G 0B1, Canada

## Abstract

Spindle positioning must be tightly regulated to ensure asymmetric cell divisions are successful. In budding yeast, spindle positioning is mediated by the asymmetric localization of microtubule +end tracking protein Kar9. Kar9 asymmetry is believed to be essential for spindle alignment. However, the temporal correlation between symmetry breaking and spindle alignment has not been measured. Here, we establish a method of quantifying Kar9 symmetry breaking and find that Kar9 asymmetry is not well coupled with early spindle alignment. We report the majority of asymmetric cells are not stably aligned. Rather, stable alignment, a state we define as perfect alignment, is correlated with Kar9 localization to the bud compartment, regardless of symmetry state. Our findings suggest that Kar9 asymmetry alone is insufficient for perfect alignment and reveal a possible role for Swe1 in regulating spindle alignment efficiency.

## Introduction

Asymmetric cell divisions are common in nature, mediating essential processes like development and regeneration. Asymmetric cell divisions occur when cells resulting from the division differ in size and/or cell fate. Crucial to the success of an asymmetric cell division is the precise regulation of spindle positioning. In this context, the spindle can be positioned off center such that the cytokinesis event produces daughters of different sizes. Additionally, cytoplasmic factors may be differentially distributed such that daughters differ in cell fate. Cortical cues are translated into spindle motions to ensure that the daughters inherit the appropriate factors. Both cases are illustrated in the first cell division of the *C. elegans* embryo. The polarity of the cortex leads to the spindle being positioned off center toward the posterior, where P-granules are also clustered^1^. The division results in a large somatic cell, and a smaller cell that contains the P-granules, which will develop into the germline of the worm^2^.

The budding yeast *S. cerevisiae* also divides asymmetrically. Growth factors must be concentrated at a prespecified bud site, to ensure emergence and growth of the bud. This requires the reorganization of the actin cytoskeleton to allow polarized transport of cargoes^3^. Specifically, the long clusters of actin filaments known as actin cables originate from the bud tip and bud neck through the localization of formins Bni1 and Bnr1 respectively^4^. This ensures that cables emanate from the bud, allowing polarized transport of cargoes via the type V myosin Myo2^3^.

As the bud neck is the future plane of cytokinesis, the spindle must position itself relative to the bud neck to ensure proper segregation of chromosomes. Moreover, the spindle itself has asymmetric properties. The nucleators of the spindle and equivalent to the centrosomes – the spindle pole bodies – undergo conservative replication, resulting in an old (pre-existing) and new (newly synthesized) spindle pole (reviewed in^5^). It is the old spindle pole that is preferentially oriented toward and inherited by the bud in 98% of cases^6^.

A unifying feature of spindle positioning is the coupling of cortical cues and spindle movements. This crucial element is mediated by SPB nucleated microtubules (MTs) that project into the cytoplasm known as astral MTs (aMTs). The aMTs exert forces on the SPBs to correctly position the spindle through their +end interactions with MT associated proteins (MAPs), MT +end tracking proteins (+TIPs), cortical proteins and motor proteins. The proteins at the cortex allow the aMTs, and by extension the SPBs, to read polarity cues and translate them into spindle pole motions leading to the alignment of the spindle axis with the polarity axis of the cell^2^.

In budding yeast, the +TIP Kar9 is responsible for aligning the spindle and orienting the old spindle pole toward the bud. Kar9 interacts with the cargo-binding domain of the type V Myosin Myo2^7^. Whether this interaction is subject to spatiotemporal regulation remains unclear. Evidence suggests that Kar9 may interact with Myo2 in both mother and bud compartments: aMT sweeping in the mother compartment has been used as evidence of Myo2-dependent transport^8^, while Myo2 has been demonstrated to be responsible for directing Kar9 to the bud tip^7^. Once bound, the Kar9-Myo2 interaction effectively translates the polarity of the cortex into spindle pole movements via the processive motion of the Kar9-Myo2 complex toward the bud tip^7^. Only once Kar9 enters the bud is the spindle expected to align along the polarity axis. This suggests a potential role for spatiotemporal regulation that could promote persistent Kar9-Myo2 interaction in the bud, thus ensuring effective spindle alignment.

Kar9 may interact with both the old and new spindle poles and their associated aMTs. If this is the case, both spindle poles may be directed toward the bud, resulting in spindle misalignment^8^. Therefore, it is believed that Kar9 symmetry breaking and accumulation to the old spindle pole is required to effectively align the spindle (Figure 1). Kar9 symmetry breaking is subject to regulation by Cdk1/Clb4. Kar9 is phosphorylated *in vivo* by the Cdk1/Clb4 complex at two sites (S197, S496). By blocking Cdk1 phosphorylation at these sites through S>A substitutions (S197A-S496A; *kar9AA*), a significant increase in Kar9 symmetry was found in the cell population^8^–10. The phosphorylation is believed to decrease Kar9 binding affinity to Bim1 on the new spindle pole and as a consequence Kar9 binds to the old spindle pole associated aMTs^8^ (Figure 1).

**Figure 1.**
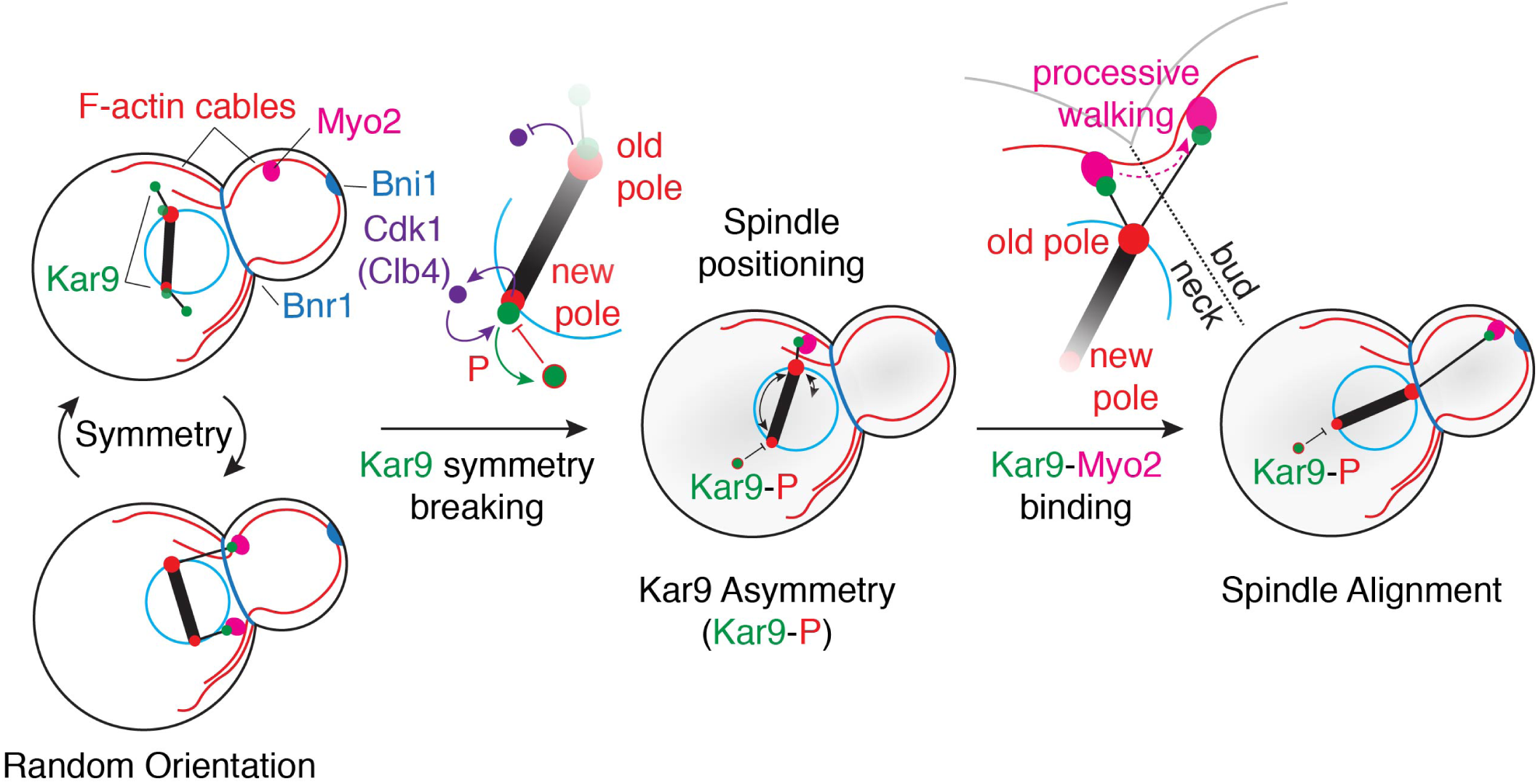
Regulation of Kar9 symmetry breaking and spindle positioning. After bipolar spindle formation, the spindle is randomly oriented. Both spindle poles and associated aMTs are decorated with Kar9. In this regime, Kar9 from both poles may interact with Myo2, thus causing both poles to be directed toward the bud, resulting in misalignment. Symmetry is broken by Cdk1/Clb4, where Kar9 at the new pole is targeted and phosphorylated by this complex. Once Kar9 asymmetry is achieved, the spindle is amenable to spindle positioning. This occurs through the processive walking of the Kar9-Myo2 complex toward the bud tip, resulting in spindle alignment.

The model shown in Figure 1 suggests that (1) symmetry breaking is an essential step to achieve spindle alignment and (2) spindle alignment occurs shortly after symmetry breaking. However, symmetry breaking and spindle alignment have not been studied as a dynamic process, therefore the temporal correlation between symmetry breaking and spindle alignment remains unclear. In the present study, we explore precisely this question by taking fast acquisitions of pre-anaphase spindles, and correlating spindle alignment with Kar9 symmetry state. We developed a method to reliably associate Kar9 foci with the spindle pole (old or new) of origin to determine symmetry states. By perturbing the symmetry breaking pathway using mutations, we were able to explore the effects of Kar9 symmetry breaking on spindle alignment. Upon examination we found that the majority of asymmetric spindles did not reach steady state alignment. Further investigation showed that steady state alignment (perfect alignment) was better correlated with Kar9 residence time in the bud rather than symmetry state, suggesting that symmetry breaking alone is insufficient to ensure spindle alignment. Moreover, our data suggests that spindle alignment requires regulation of Kar9 residence within the bud.

## Results

### A trajectory-based method of Kar9-pole association

Previous determination of Kar9 symmetry required aMT labelling to visually assign the symmetry state of each cell^8^. However, Kar9 symmetry state can be determined in the absence of aMT contouring using Kar9 trajectories by employing +TIP tracking to trace aMT dynamics at short timescales. Specifically, two consecutive Kar9 spots in a track can define a line in 3D tracing the motion of the dynamic aMT +end, eliminating the need for aMT labelling. This method relies on the fact that over the course of a short time step (e.g. 5 seconds), the spindle poles move slowly in comparison to highly dynamic aMT +ends and assumes the majority of aMTs contours are linear in this regime. Therefore, two consecutive Kar9 spots should trace the position of MT+end in relation to the spindle pole of origin that has remained nearly stationary in comparison. Since this method requires high temporal resolution, only a snapshot of the pre-anaphase spindle alignment process is captured, which normally takes 20-30 minutes to achieve. Instead of tracing symmetry breaking over time, symmetry states over the course of spindle alignment are captured and correlated with alignment of the spindle axis to the polarity axis as well as spindle length (distance between the old and new spindle poles).

Analysis methods are detailed in the Methods section. In brief, yeast expressing Kar9-mNeonGreen and SPB reporter Spc42-mKate2 were imaged every 5 seconds for 5 minutes. Brightfield images were used to establish the axes of the cell (Figure 2a). Fluorescent images were used to track objects in three-dimensions (3D) with high precision by fitting signal intensity to a 3D gaussian (implemented with TrackMate^11^) (Figure 2b).Spindle pole tracks were used to determine length and orientation of the spindle. Kar9 tracks were used to determine the pole of origin. Consecutive coordinates in the track were used to define a line in 3D, and the pole closest to the line was recorded. The operation was repeated for every consecutive pair, and the pole closest to the track over 80% of the time was deemed the pole of origin (Figure 1c,d). Symmetry was then determined based on Kar9-pole association. Spindles were classified as asymmetric when all Kar9 tracks pointed to a single pole (Figure 2e,f). Conversely, spindles were classified as symmetric when a fraction of the tracks pointed to the old pole and others the new pole (Figure 2g-j).

**Figure 2.**
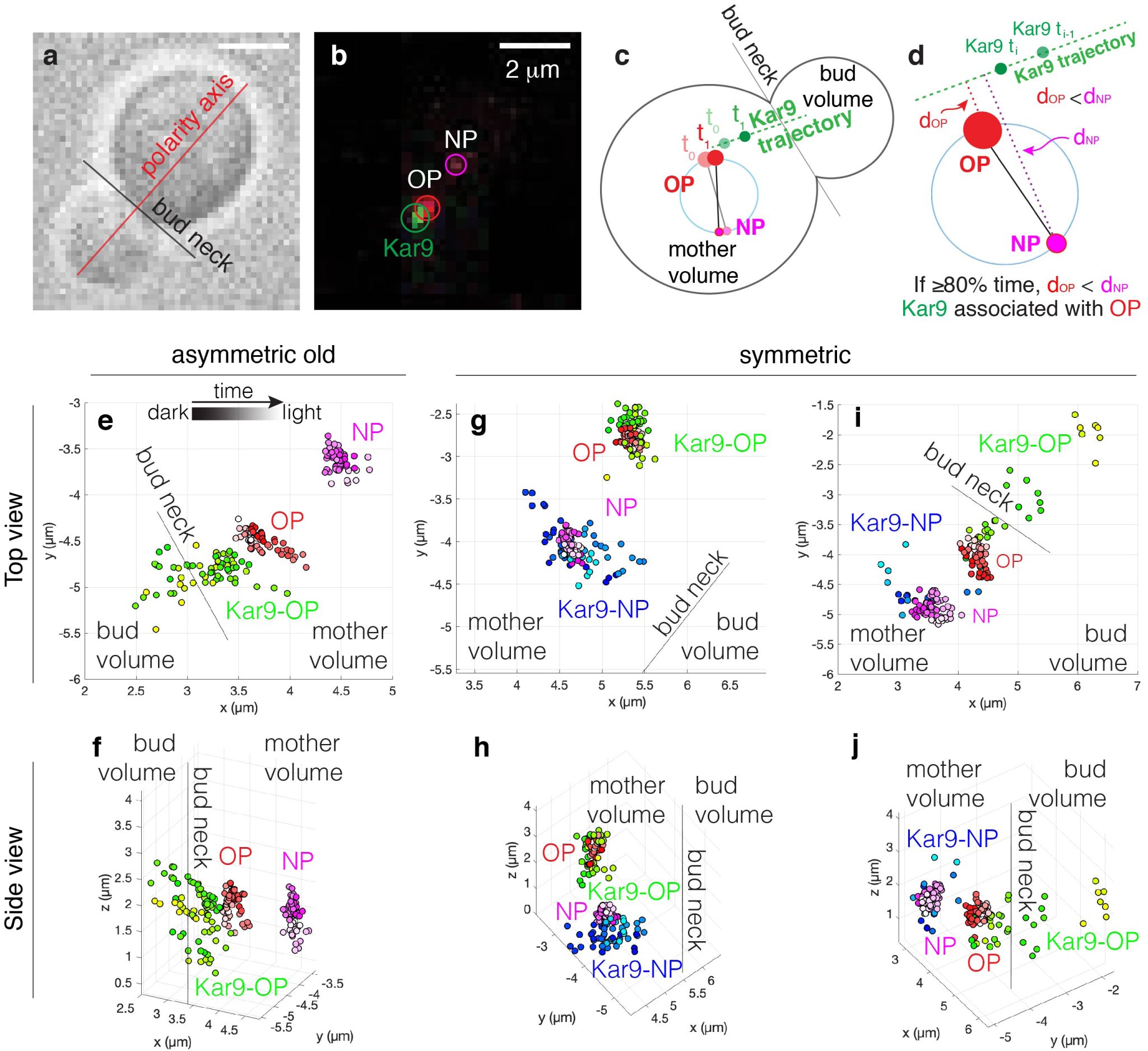
A trajectory-based method of Kar9-pole association. (a) Representative brightfield image used to determine bud neck and polarity axis. (b) Representative fluorescent image (max-projected). Old and new pole are differentiated based on fluorescence intensity. Scale bar 2 *µ*m. (c-d) Graphical description of trajectory method. (e-f) An example of an asymmetric old state: Kar9 clearly associates with the old pole exclusively. (g-h) An example of a symmetric state in which Kar9 associates with both poles over the entire acquisition. (i-j) An example of a cell that fluctuates between symmetry and asymmetry which in my study is classified as symmetric. Each representative cell is shown from a top view (e,g,i) and side view (f,h,j).

Upon examination of wild type (WT) cells, we found a large heterogeneity in symmetry states. Kar9 exhibited a variety of behaviours, such as exclusive association with the old or new pole (asymmetric old or asymmetric new respectively, Figure 2e,f) or association with both poles over the entire acquisition (symmetric, Figure 2g,h). Furthermore, a fraction (21 of 132) spindles were observed to fluctuate between symmetry and asymmetry. In such cases, we did not consider symmetry to have irreversibly broken; any cell that displayed such fluctuations between states was classified as symmetric (Figure 2i,j). To validate the method used in the present study, frequencies of Kar9 symmetry and asymmetry in WT and *kar9AA* populations were examined and compared to previously published work. As *kar9AA* cells have previously been characterized to be overwhelmingly symmetric^8^, classification of the population through our method was expected to yield a similar frequency of symmetry. Indeed, the *kar9AA* population was primarily symmetric (73%, 96 of 131 cells, Figure 3b, inset) in comparison with the WT population which was primarily asymmetric (59%, 78 of 132 cells, Figure 3a, inset).

**Figure 3.**
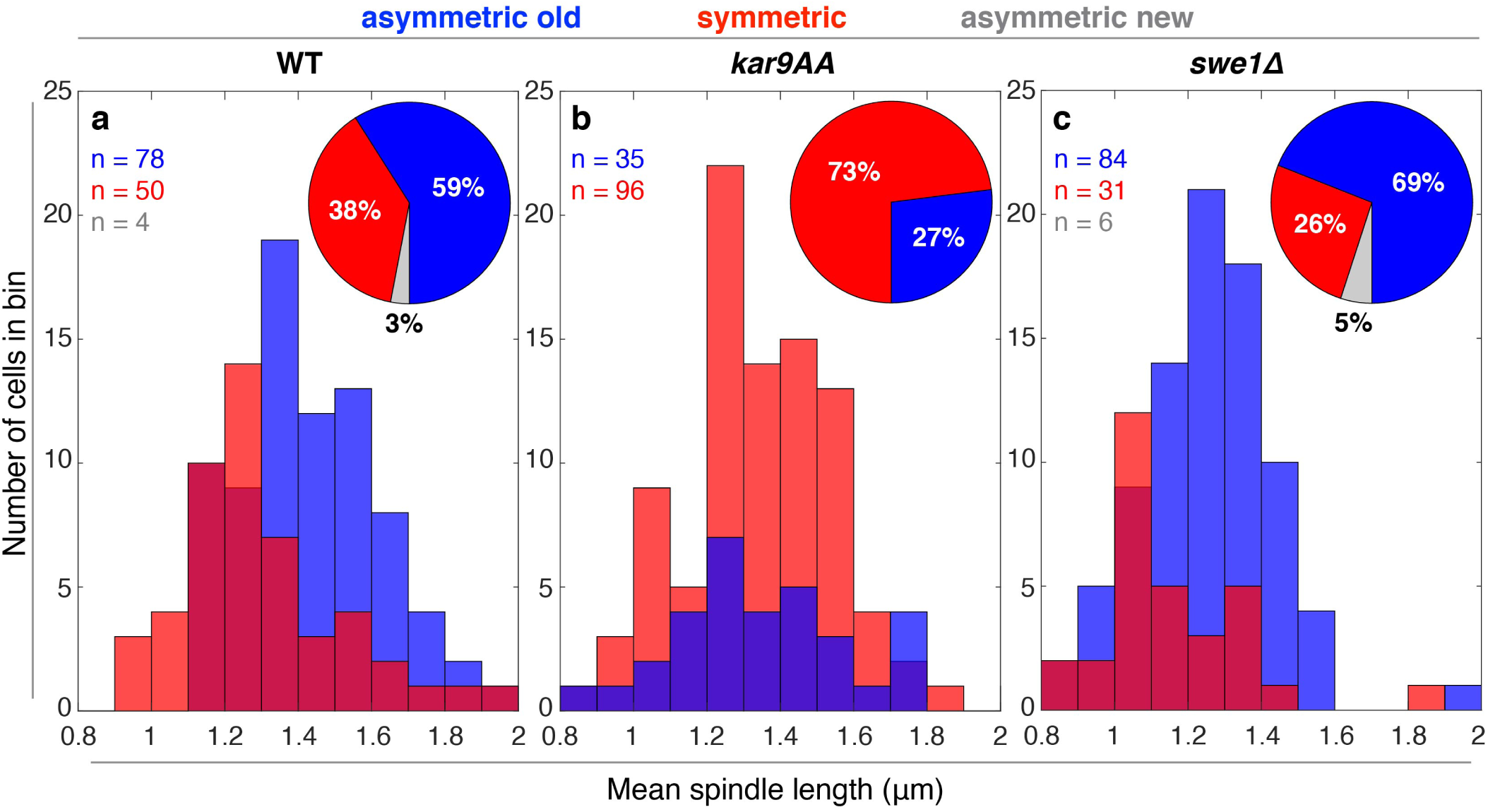
Genetic perturbations affect the timing of symmetry breaking. (a-c, inset) Proportion of asymmetric old, symmetric and asymmetric new spindles in WT(a), *kar9AA*(b), and *swe1*Δ(c) populations. (b,inset) Blocking Cdk1 regulation of Kar9 effectively increases Kar9 symmetry in the population. (c, inset) *swe1*Δ cells display increased asymmetry in the population. (a-c) Spindle length distributions. (a) Symmetry breaking is correlated with increasing spindle length, and approach to anaphase in WT cells. (b) Symmetry breaking occurs at random in the *kar9AA* population. (c) Deletion of Swe1 results in early symmetry breaking.

### Genetic perturbations affect symmetry breaking efficiency

Next, we sought to examine the timing of symmetry breaking. Absolute time spent is difficult to measure, as each cell progresses through the cell cycle at different rates. However, the pre-anaphase bipolar spindle grows at a relatively constant rate of 0.15 nm/s in WT cells and mutants used in this study (Supplementary Figure S1), therefore mean spindle length can be used as a proxy for time. Short spindles (0.8 *µ*m) indicate early bipolar spindle formation and increasing spindle length is proportional to increasing time spent in metaphase. By correlating spindle length with symmetry state, we report that the majority of symmetric WT spindles were short, with a sharp drop at about 1.3 *µ*m in length. The majority of asymmetric WT spindles were found at relatively longer spindle lengths (Figure 3a). Therefore, symmetry is correlated with early spindle formation, and symmetry breaking is correlated with increasing time.

In the absence of Cdk1 regulation of Kar9 via the *kar9AA* mutation, symmetry breaking is expected to be uncoupled from progression to the metaphase/anaphase transition, with symmetry breaking occurring at random. Indeed, our analysis revealed symmetry state to be uncoupled from spindle length. Instead, symmetric and asymmetric populations were similar: they were broadly distributed along the entire range of short spindle lengths, indicating a lack of regulation in the system (Figure 3b).

We next decoupled Cdk1 inhibition of Kar9 from cell cycle progression by deleting the Wee1 ortholog Swe1^12^. Swe1 inhibits both early (Clb3,4) and late (Clb1,2) M-phase forms of Cdk1 in budding yeast^13,14^. Therefore, M-phase Cdk1 activity is expected to be unrestrained in the *swe1*Δ mutant with symmetry breaking occurring earlier than in WT cells. *swe1*Δ cells spanned a smaller range of pre-anaphase spindle lengths, consistent with the role of Swe1 in mediating the metaphase-to-anaphase transition^15^. The difference in spindle length may also be attributed to the smaller size of the cells. As expected, we observed an increase in the number of asymmetric cells in the population (69%, 84 of 121 cells, Figure 3c, inset). Importantly, we observed a subset of Kar9 asymmetric cells in the population at short spindle lengths that was not observed in WT asymmetric cells. This suggests that symmetry breaking occurs earlier in this condition (Figure 3c).

### Perturbations in the symmetry breaking pathway alter alignment efficiency

By employing *kar9AA* and *swe1*Δ mutants, we are able to examine how spindle alignment responds to a decrease or increase in symmetry breaking respectively. To characterize the alignment of different symmetry states, we employed a previously developed method to characterize 3D spindle alignment^16^. The 3D spindle coordinates are projected onto the polarity axis, resulting in a one-dimensional (1D) projected length. The 1D length is plotted versus the true 3D length to determine alignment. The spindle is aligned when 1D and 3D lengths are comparable. To simplify the analysis, we defined an alignment index (AI) as the mean ratio of 1D to 3D length for each spindle. A numerical value of AI *≈* 1 indicates proper alignment and orientation (old pole proximal to the bud). A numerical value of AI *≈* -1 implies proper alignment but misorientation (new pole proximal the bud). All other AI values ranging from -1 to 1 designates various degrees of misalignment, where an AI *≈* 0 indicates the spindle is perpendicular to the polarity axis. Regardless of genotype, the majority of symmetric spindles were broadly distributed about an AI value of 0, indicating poor alignment (Figure 4a-c). This is consistent with a regime in which both poles are directed to the bud tip (Figure 1). WT asymmetric spindles were biased toward a high AI (Figure 4a). *kar9AA* asymmetric spindles showed defects in alignment, despite their asymmetry (Figure 4b). Remarkably, while *swe1*Δ cells break symmetry earlier, the asymmetric spindles were less efficient in alignment relative to their WT counterparts (Figure 4c). All 3D spindle measures are consistent with 2D alignment measures (Supplementary Figure S2).

**Figure 4.**
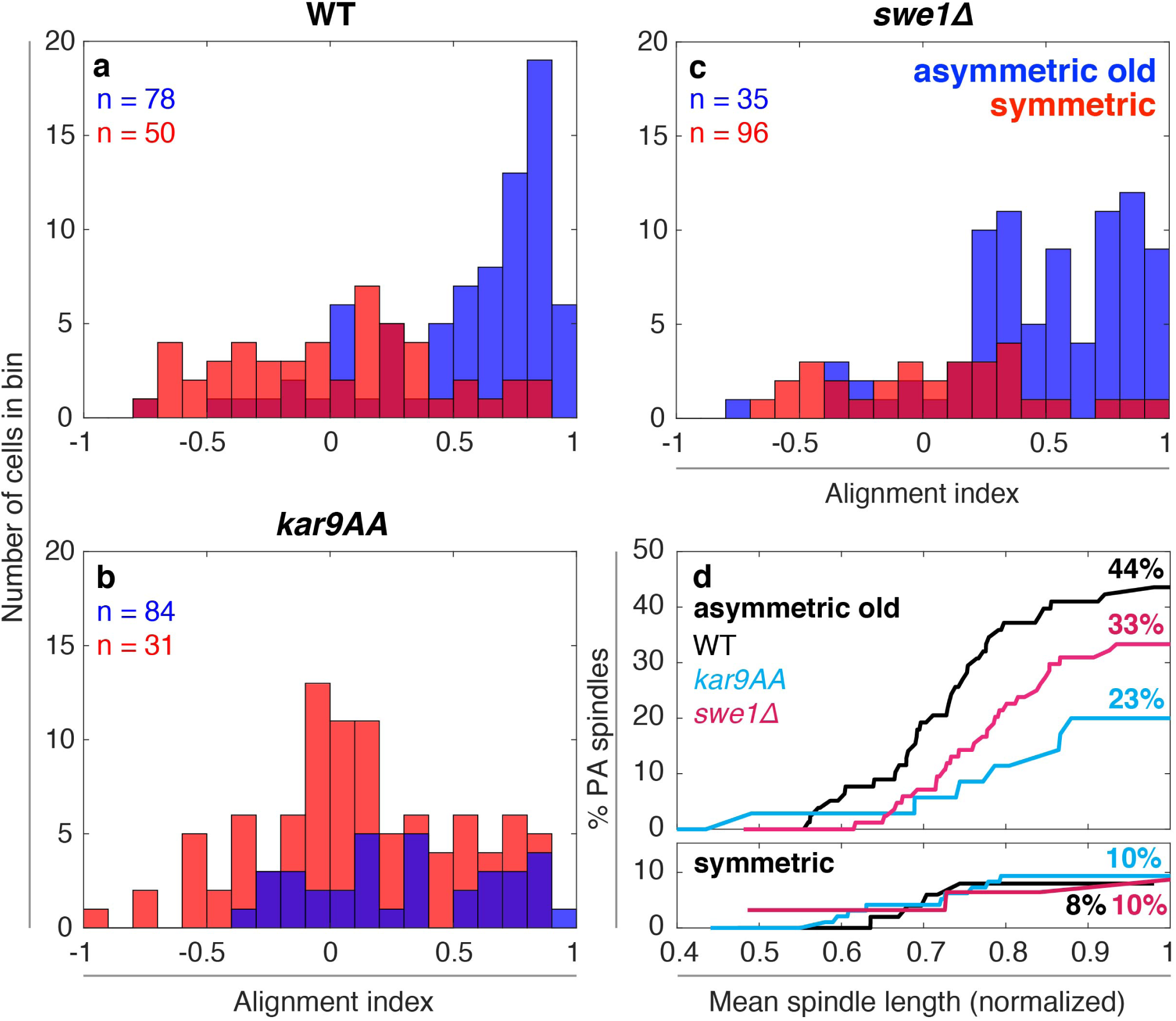
Altering symmetry in a population does not proportionally alter alignment efficiency. (a-c) Spindle alignment of WT(a), *kar9AA*(b) and *swe1*Δ(c) populations. (a) WT asymmetric cells are heavily biased toward better alignment (AI approaching 1). (b) *kar9AA* asymmetric spindles are defective in alignment. (c) *swe1*Δ asymmetric spindles are biased toward better alignment but are less efficient then WT cells. All symmetric cells regardless of genotype are more broadly and uniformly distributed across the range of AIs. (d) Cumulative percentage of PA spindles in asymmetric (top) and symmetric (bottom) populations. Asymmetric spindles are able to achieve perfect alignment more efficiently that symmetric cells, but efficiency is decreased in the *swe1*Δ and even more so in the *kar9AA*. Symmetric spindles regardless of genotype achieve perfect alignment at similar rates.

In order to better characterize spindle alignment in these populations, we classified spindles as perfectly aligned (PA) or not aligned (NA) based on alignment index. If the absolute value of the AI is greater than 0.75, the spindles are perfectly aligned (AI *≥* 0.75, PA toward the old pole; AI *≤* -0.75, PA toward the new pole). All other spindles are classified as NA. This threshold represents the quality of the spindle’s alignment. If the spindle is oriented such that one pole is able to enter the bud if the cell were to undergo anaphase at that point, the spindle is PA. However, if the orientation of the spindle does not allow for SPB entry upon anaphase, the spindle is classified as NA.

Using these categorizations, we unexpectedly found that the majority of asymmetric spindles were not perfectly aligned (44%, 34 of 78 cells, Figure 4d), suggesting that perfect alignment is poorly correlated with asymmetry. Furthermore, perturbations in the symmetry breaking pathway decreased the frequency of PA spindles. This is especially notable in the *swe1*Δ asymmetric population, where only 33% of spindles were PA, despite an overall increase in the frequency of asymmetric spindles (28 of 84 cells, Figure 4d). Moreover, there were a subset of symmetric spindles that were perfectly aligned, supporting the observed poor correlation between symmetry breaking and spindle alignment (Figure 4d).

The 5-minute acquisition length does not allow for the observation of the entire spindle alignment process. However, we can estimate when perfect alignment is achieved in the population by using spindle length as a proxy. Since *swe1*Δ spindles undergo the metaphase-to-anaphase transition at shorter spindle lengths^15^, spindle length is normalized to the mean spindle length prior to the onset of anaphase. To ensure spindle growth was comparable between strains, we measured the mean spindle growth over the 5-minute acquisition for each population. Indeed, all three strains demonstrated similar growth distributions (with a mean of 0.15, 0.26 and 0.09 nm/s for WT, *kar9AA* and *swe1*Δ respectively). Furthermore, the computed spindle growth was independent of mean or initial spindle length (Supplementary Figure S1).

WT asymmetric spindles steadily became more perfectly aligned as spindle length increased, while WT symmetric ones did not (Figure 4d). A similar trend was observed in the *kar9AA* population. However, fewer asymmetric *kar9AA* cells achieved perfect alignment at a given spindle length relative to WT asymmetric cells (Figure 4d). Asymmetric spindles in the *swe1*Δ population also achieved perfect alignment as metaphase progressed. While more efficient than *kar9AA* cells, *swe1*Δ asymmetric cells achieved perfect alignment less efficiently than the WT (Figure 4d). Altogether, our data suggests a disparity between symmetry breaking and alignment: while cells are asymmetric, the majority of spindles are not aligned. Furthermore, increasing and decreasing rates of symmetry breaking in the population via the *swe1*Δ and *kar9AA* mutants do not proportionately affect alignment, rather they are both detrimental to reaching perfect alignment.

### Kar9 residence in the bud predicts alignment efficiency

Evidence clearly suggests that perfect alignment and asymmetry are not well correlated, prompting us to investigate what else was needed to achieve perfect alignment. Therefore, we examined Kar9 localization to the mother and bud compartments. Quantifying Kar9 localization to the bud revealed that WT-PA spindles displayed increased Kar9 residence time in the bud relative to that of NA spindles, regardless of symmetry state (Figure 5a). This was also true in strains with mutations in the symmetry breaking pathway (Figure 5b,c), suggesting Kar9 residence in the bud to be an important aspect of spindle alignment independent of symmetry breaking. Furthermore, the extent of Kar9 residence time in the bud was consistent with alignment trends of WT, *kar9AA* and *swe1*Δ populations. While perfect alignment is correlated with time spent in the bud, Kar9 of *kar9AA* and *swe1*Δ populations spends less time in the bud relative to their WT counterparts, which may explain why these populations are less effective at spindle alignment.

**Figure 5.**
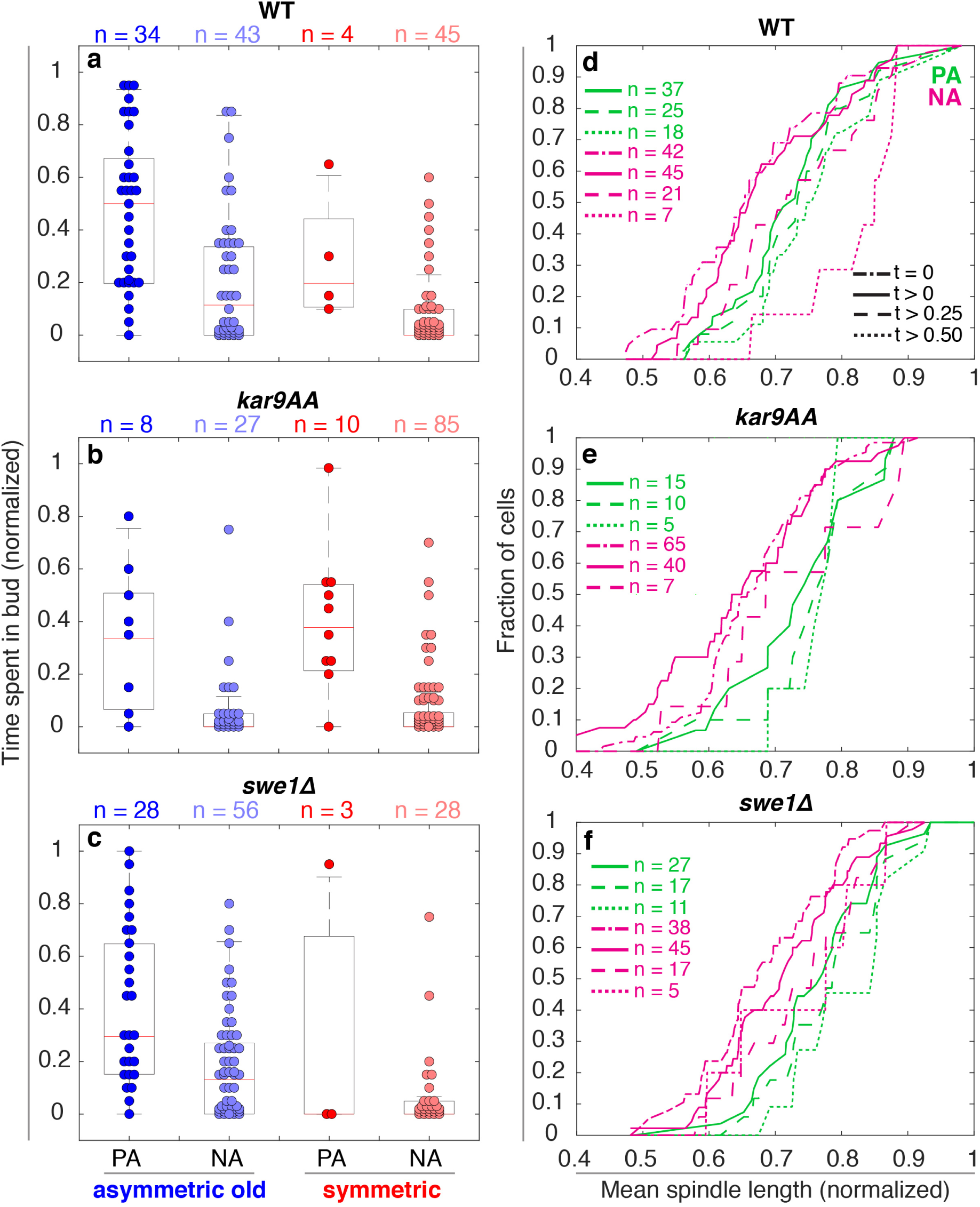
Bud entry and time spent in bud predicts alignment efficiency. (a-c) Number of time points spent in bud normalized to length of the acquisition. (a) Kar9 of PA-asymmetric spindles spends more time in the bud than that of NA-asymmetric spindles. The same is true for PA and NA symmetric spindles. (b-c) *kar9AA* and *swe1*Δ population display a similar pattern to WT cells, but overall spend less time in the bud. (d-f) Cumulative number of cells with Kar9 residence times above an indicated threshold. (d) The number of WT spindles displaying non-zero residence times increases with spindle length (e) Similarly, residence time in the bud is correlated with metaphase progression in *kar9AA* cells. (f) *swe1*Δ consistently display delays in the accumulation of non-zero residence times, irrespective of the state of alignment.

To further examine a potential spatial requirement for alignment, we measured the extent to which Kar9 explored the bud in PA and NA spindles, irrespective of symmetry state. The 3D position of Kar9 was projected onto the polarity axis, and the depth was defined as the distance between the projected coordinate and the midpoint of the bud neck. Values smaller than 0 indicated localization to the mother compartment, while those above 0 localization to the bud (Supplementary Figure S3). Depth was normalized to accommodate for variance in bud size. Therefore, a Kar9 depth of 1 indicates Kar9 presence at the bud tip.

Regardless of symmetry state or genotype, Kar9 depth of PA spindles remained exclusively in the bud and region surrounding the bud neck (between -0.5 and 0.5, Supplementary Figure S3). NA spindles spanned a much broader range of depths and rarely resided in the bud and bud neck region, consistent with measures of time spent in the bud (Supplementary Figure S3). Kar9 of *kar9AA* and *swe1*Δ cells much less frequently localized deeper within the bud, consistent with the observed drop in Kar9 residence times and inefficiency in spindle alignment (Supplementary Figure S3).

We inquired whether Kar9 depth and residence time in the bud was correlated with alignment efficiency. Indeed, as Kar9 entered deeper into the bud and resided longer, the AI of WT, *kar9AA* and *swe1*Δ spindles approached 1 (approach PA, Supplementary Figure S3). Importantly, this trend persisted regardless of symmetry state, further suggesting that achieving asymmetry does not ensure alignment.

Next, we asked whether Kar9 but entry was correlated with metaphase progression, irrespective of symmetry state. To do so, we pooled cells into PA and NA classes, and examined the cumulative number of cells with Kar9 residence times equal to or above a given threshold (t = 0, t *>* 0, 0.25, 0.50, Figure 5d-f). Spindle length was normalized to the mean spindle length prior to anaphase onset^15^ to facilitate comparison between strains. Groups with a sample size less than 5 cells were omitted for clarity, but are displayed in Supplementary Figure S4.

At a normalized length of ≈ 0.55 (1.1 *µ*m), WT PA spindles with non-zero residence times began to accumulate. Those spindles displaying residence times over 0.25 and 0.50 followed shortly at similar rates. The large majority of PA spindles with non-zero residence times spent over half of the time in the bud (51%, 18 of 37 cells, Figure 5d). Conversely, WT NA spindles of zero and non-zero residence times accumulated similarly as metaphase progressed. Less than half (46%, 21 of 45) resided in the bud for a quarter of the acquisition, and only began to accumulate later in metaphase (normalized length of *≈* 0.55, 1.1 *µ*m, similar to PA spindles). Unlike PA spindles, few NA spindles displayed residence times exceeding half the acquisition. These events largely occurred late in metaphase (normalized length *≈* 0.85, 1.7 *µ*m) (Figure 5d). Notably, at a normalized length of *≈* 0.7 (1.4 *µ*m), the number of PA spindles with non-zero residence times began to accumulate more rapidly. Meanwhile, NA spindles with non-zero residence times became more infrequent at the normalized length of *≈* 0.75 (1.5 *µ*m) (Figure 5d). This suggests a potential transition from random Kar9 bud entry to Kar9 retention within the bud, leading to perfect alignment.

Similarly, *kar9AA* PA spindles with non-zero residence times began to accumulate at a normalized length of *≈* 0.5 (1 *µ*m). In contrast with the WT, only one third of *kar9AA* PA spindles displayed residence times exceeding half of the acquisition (5 of 15). These were also delayed with respect to the WT, beginning their accumulation at a normalized length of *≈* 0.7 (1.4 *µ*m, relative to 0.55 in WT PA cells) (Figure 5e). *kar9AA* NA spindles exhibited similar patterns to that of WT NA spindles; shifted similarly in time and accumulating earlier than *kar9AA* PA spindles (Figure 5e).

As *swe1*Δ cells were less efficient in alignment, we asked whether the mutant influenced alignment efficiency by altering Kar9 retention within the bud. *swe1*Δ PA spindles with non-zero residence times began accumulating at a normalized length of *≈* 0.65 (1.11 *µ*m), delayed relative to the WT which began its accumulation at a normalized length of 0.55. The distance between residence time thresholds in PA spindles was accentuated in *swe1*Δ cells relative to the WT (Figure 5f). Furthermore, zero and non-zero residence times in NA spindles of the other two strains accumulated concurrently, however *swe1*Δ NA spindles were delayed in achieving non-zero residence times (Figure 5f). Altogether, the consistent shifts toward longer normalized lengths observed in *swe1*Δ cells suggests Swe1 may promote the retention of Kar9 in the bud, thus facilitating spindle alignment.

## Discussion

In this study, we investigated the extent to which Kar9 symmetry breaking and efficient alignment are temporally correlated. By developing a method to dynamically assay Kar9 symmetry breaking, we were able to scrutinize its relation to spindle alignment as a function of spindle length. The spindle elongates at a relatively constant rate of 0.15 nm/s during metaphase, therefore spindle length serves as a proxy for time. We were surprised to find a poor correlation between symmetry breaking and efficient spindle alignment (perfect alignment). The low number of WT asymmetric cells displaying perfect alignment suggests that symmetry breaking does not rapidly align the spindle (Figure 4). To further examine the coupling between Kar9 symmetry breaking and spindle alignment, we manipulated the timing of Kar9 symmetry breaking using genetic perturbations known to inhibit Kar9 asymmetry (*kar9AA*) or increase Cdk1 activity in metaphase (*swe1*Δ). While frequencies of symmetry/asymmetry in mutant populations were altered by the imposed genetic perturbations, frequencies of perfect alignment were not proportionally affected. Notably, *swe1*Δ cells broke symmetry earlier yet produced fewer perfectly aligned spindles; such cells were delayed in achieving perfect alignment. Furthermore, Kar9 localization and persistence in the bud compartment was correlated with stable “perfect” spindle alignment. This strongly suggests that Kar9 retention within the bud is far more important in regulating the timing of spindle alignment than symmetry breaking itself. By examining the effects of *kar9AA* and *swe1*Δ on Kar9-bud localization, we identified delayed retention of Kar9 in *swe1*Δ cells, thus highlighting Swe1 as a potential mediator of Kar9-bud localization and spindle alignment.

The consequences of the imposed perturbations on the symmetry breaking pathway suggest that symmetry breaking and alignment are partially independently regulated. The number of PA spindles in the WT asymmetric class may simply be an indication of the inherent efficiency of the system: reaching alignment takes time, therefore only a subset of cells in the unsynchronized population achieved it. If alignment is a consequence of symmetry breaking, and not an independent process, frequencies of perfect alignment in asymmetric populations should be comparable, regardless of genotype. The *kar9AA* and *swe1*Δ mutants altered symmetry distributions, making a population more or less symmetric respectively. However, asymmetric *kar9AA* and *swe1*Δ cells both displayed decreased alignment efficiency with respect to the WT. This suggests that the mutants influence alignment in addition to and independently of symmetry breaking, and the poor correlation between symmetry breaking and alignment is due to differential regulation of the two processes.

Furthermore, spindle alignment may be subject to spatiotemporal regulation. Kar9 residence time in the bud was correlated with perfect alignment, regardless of symmetry state (Figure 5). This highlights bud entry as an important parameter in spindle alignment. The ability to retain Kar9 in the bud compartment was correlated with metaphase progression, occurring more efficiently after a normalized length of *≈* 0.75 (1.5 *µ*m) in WT cells. While our data implies spatiotemporal regulation of spindle alignment, further quantification of Kar9-Myo2 interactions in space and time is required. Moreover, our data suggests that Swe1 mediates the efficiency of Kar9 retention in the bud. We observed a decrease in the number of PA spindles displaying large (t > 0.5) residence times in the *swe1*Δ population. These cells accumulated later in metaphase relative to the WT. We speculate that Swe1 may be able to influence spindle alignment by mediating Kar9-Myo2 interactions. Swe1 may post-translationally modify Kar9 and/or Myo2 to increase their affinity to one another, thus retaining Kar9 in the bud, effectively aligning the spindle.

By employing spatiotemporal regulation of alignment, the cell is able to diminish instances of spindle misalignment. Assuming persistent Kar9-Myo2 attachments are not restricted in space or time, more symmetric spindles should display misalignment (AI *≈* 0). However, WT symmetric spindles are uniformly distributed in orientation. Only when the temporal separation between symmetry breaking and alignment is disrupted in the *kar9AA* population do we begin to see an accumulation of misaligned symmetric spindles (Figure 4b). Moreover, the likelihood that aMTs from both spindle poles enter the bud is low, thus by imposing this spatial requirement on alignment the cell further ensures that only one spindle pole is directed toward the bud.

Altogether, we propose an alternate model to the pre-anaphase spindle alignment process, in which spatiotemporal regulation of symmetry breaking and spindle alignment is incorporated. Symmetry breaking and alignment occur in distinct compartments: symmetry breaking in the mother compartment, and alignment in the bud compartment (Figure 6b). Symmetry breaking, facilitated by Cdk1/Clb4 early in metaphase (*≈* 1.3*µ*m), allows the spindle to be amenable to meaningful spindle movements that will ultimately result in spindle alignment (Figure 6). However, the spindle will not be aligned without persistent Kar9 localization in the bud, where Kar9-Myo2 interactions are potentially strengthened by Swe1 later in metaphase (*≈* 1.5*µ*m) (Figure 6). Due to the compartmentalization of these two processes, they likely occur partially independently: achieving asymmetry does not ensure spindle alignment, and perfect alignment can be observed in cells with Kar9 symmetry.

**Figure 6.**
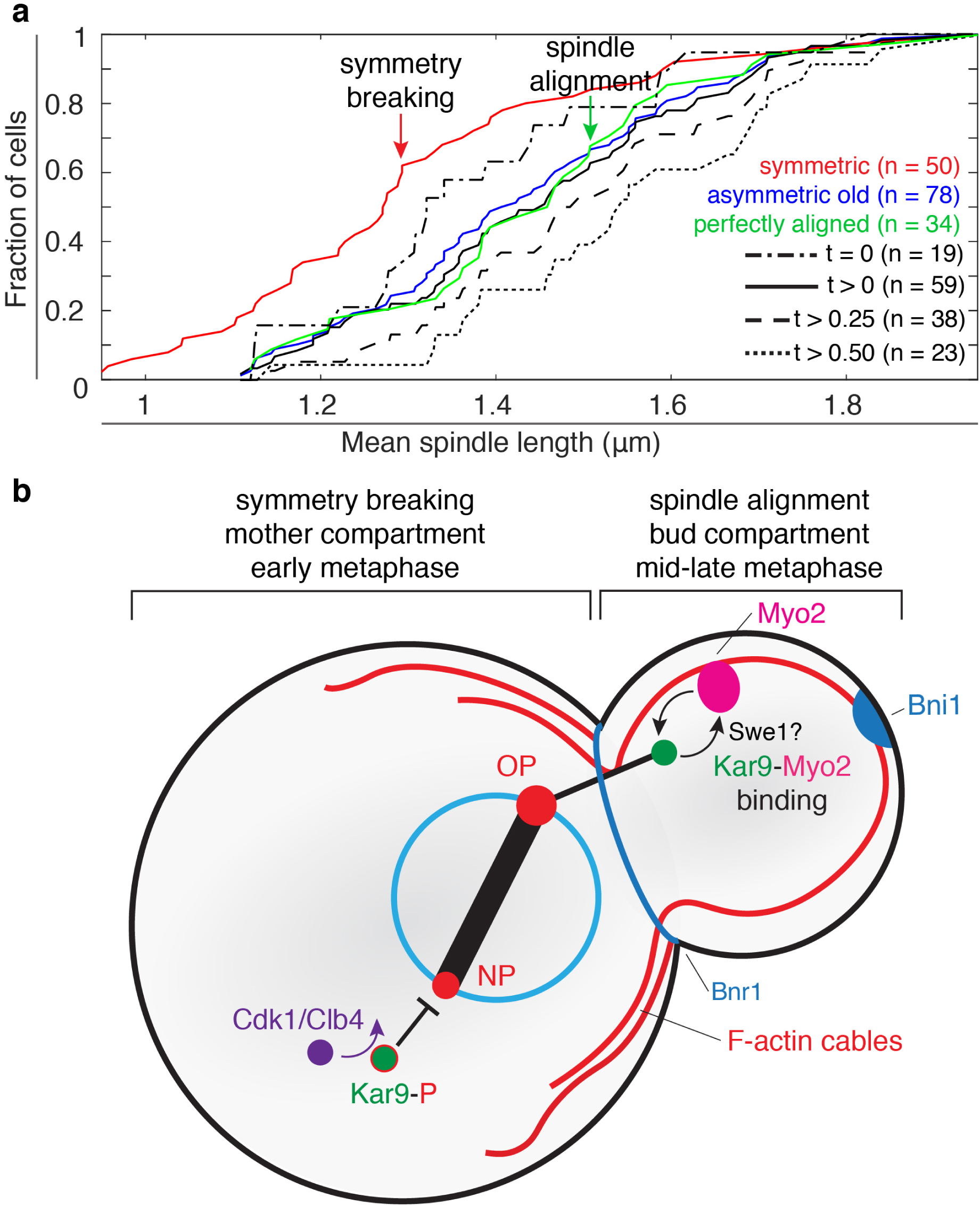
Symmetry breaking and spindle alignment are spatially and temporally separated. (a) In WT cells, symmetry breaking occurs early in metaphase, largely occurring prior to reaching a spindle length of 1.3 *µ*m. Of the cells displaying Kar9 asymmetry, the likelihood of bud entry and retention within the bud increases as metaphase progresses. At mid-to-late anaphase, asymmetric spindles begin to achieve perfect alignment. (b) Early in metaphase, Kar9 symmetry is broken by Cdk1/Clb4 in the mother compartment. Later in metaphase, Kar9-Myo2 interactions may be modified (possibly through Swe1) to promote attachment, resulting in longer residence times in the bud, ultimately leading to spindle alignment.

It is interesting to contemplate the potential role of mitotic exit network (MEN) kinases in the regulation of spindle alignment. Asymmetric *kar9AA* cells were the most deficient in effective alignment (Figure 4), suggesting the amino acid substitutions may alter Kar9’s functionality. Notably, MEN kinases Dbf2/20 were shown to target Kar9, including S197, which is blocked by the *kar9AA* mutation^10^. Therefore, MEN kinases may be involved in Kar9-mediated spindle alignment. Spindle positioning and the MEN are coupled by the localization of Tem1 activators and inhibitors in the bud and mother compartment respectively. This differential localization of activators and inhibitors creates activation zone in the bud and inhibitory zone in the mother^17^. Thus the position of the SPBs act as a sensor, as Tem1 remains inactive in the mother and only becomes active upon bud entry^17^. Due to the nature of the relationship between spindle positioning and the MEN, and Kar9’s role in positioning the spindle proximal to the bud neck, it is possible that the MEN contributes to spindle alignment through Kar9.

## Methods

### Strain construction

All strains used are derived from BY4741 and are listed in Supplementary Table S1. All fluorescently tagged proteins are expressed under their endogenous promoter.

### Growth conditions and live cell imaging

Yeast was grown overnight at 25*°*C in 5mL synthetic complete (SC). Liquid cultures were diluted to 0.2 OD and incubated 1.5-2 hrs at 25*°*C to reach log phase before imaging. Imaging was performed at 25*°*C on a custom built dual-camera spinning disk confocal microscope with the following components: a Leica DMi8 inverted microscope with Quorum Diskovery platform installed, 63x/1.47 numerical aperture objective, 50 *µ*m pinhole spinning disk, two iXon Ultra 512×512 EMCCD cameras for synchronous acquisitions, 488 nm and 561 nm solid state lasers, and an MCL nano-view piezo stage with ASI three axis motorized stage controller. Imaging was mediated through MetaMorph acquisition software. 488 nm and 561 nm lasers at 25% and 35% power respectively were used to excite fluorophores simultaneously. 14 z-stacks of 300 nm step size and 55 ms exposure were taken every 5 seconds for a total of 5 minutes. Data is pooled from 3-4 independent experiments for each strain.

### Segmentation and tracking

Individual cells were manually segmented using BF images. The bud neck coordinates were determined by recording the coordinates of the inflection points of the bud neck. The polarity axis was determined as the axis perpendicular to the bud neck. The angle between the horizontal and the polarity axis was measured (0 to 360*°*): angle ranged from 0 to 180*°* if the bud faced upward and 180*°* to 360*°* if the bud faced downward. Bud size was measured as the distance between the bud tip and the midpoint of the bud neck. All measurements were taken using the basic Fiji toolbox.

Fluorescent images were tracked with high precision in 3D using Fiji plugin TrackMate^11^. A difference of gaussian filter (DoG) was used to detect fluorescent spots. A threshold intensity of 200 and 270 was used to detect mKate2 and mNeonGreen objects respectively. The simple Linear Assignment Problem (LAP) algorithm was used to link spots into tracks^18^. A distance threshold of 1 *µ*m and 1.5 *µ*m was input for Spc42 and Kar9 respectively. Gap-closing was allowed for Spc42 (max distance 1 *µ*m, max length 2 frames), but not for Kar9. Track coordinates were exported for analysis in MATLAB.

### Kar9-pole association algorithm

A Kar9 trajectory vector was defined from the position of two consecutive Kar9 objects:

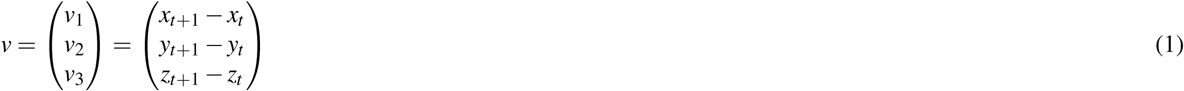

The vector was used to define a line tracing the trajectory of the motion:

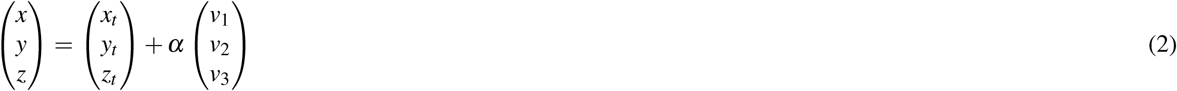

The shortest distance between the line and each pole (*x*_*p*_, *y*_*p*_, *z*_*p*_) is perpendicular to the trajectory line. Therefore, the dot product of the two vectors must be zero:

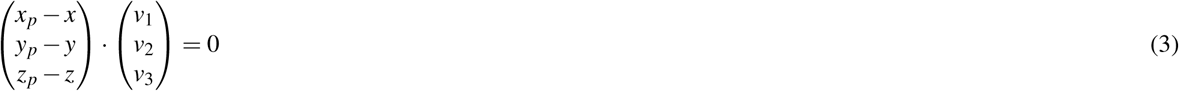

Expanding and substituting *x, y* and *z* for their respective values:

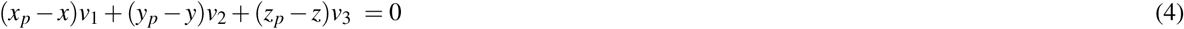

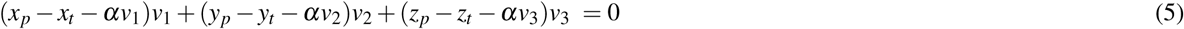

Solving for the unknown variable *α*:

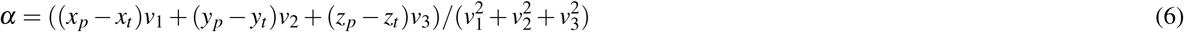

The obtained value of *α* is used to determine the distance between each pole and the trajectory line:

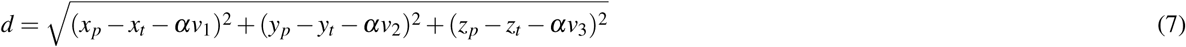

The closest pole is recorded. The measurement is repeated for each consecutive pair of Kar9 tracks. The pole most often proximal to the respective trajectory (80% of the time or more) is deemed the pole of origin. Asymmetric old (new) cells have all tracks associated with the old (new) pole. Symmetric cells have at least one track associated with the old pole and at least one track associated with the new pole. Tracks below the 80% threshold are discarded. Tracks of an entire cell are discarded if a track with less than 80% linkage interferes with ability to determine the symmetry state of the cell.

### 2D alignment measures

A vector defined by the coordinates of the spindle poles is defined:

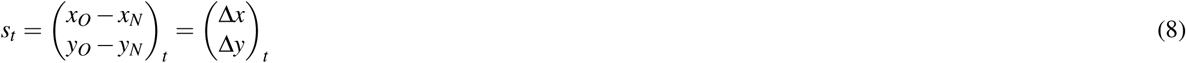

The angle (*θ*_*t*_) between the horizontal and the vector *s*_*t*_ is determined:

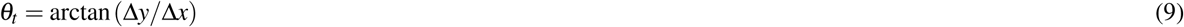

Finally, the absolute difference between *θ*_*t*_ and the angle defined by the polarity axis (*ϕ*) is calculated:

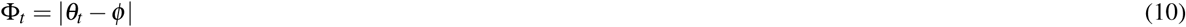

This calculation is performed and recorded for every time point in the acquisition.

### 3D alignment measures

This method is adapted from Shulist et al., 2017^16^. A vector defined by the polarity axis is determined using the measured polarity axis angle (*ϕ*):

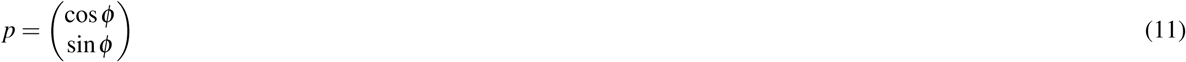

The 3D SPB coordinates are first projected onto the XY plane simply by retaining only *X* and *Y* coordinates. They are then projected onto the polarity axis vector p. The magnitude of the projected, 1D vector is determined:

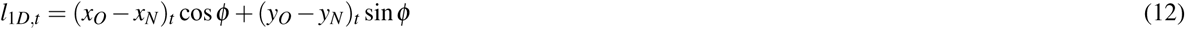

The calculation is repeated for each time point t in the acquisition. The alignment index is defined as the mean ratio of 1D to 3D lengths for the entire acquisition:

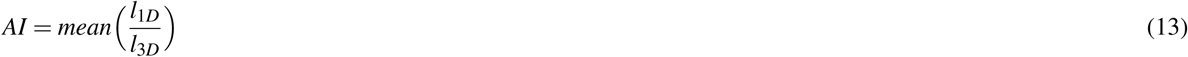

### Spindle length

The spindle length is simply determined as the distance between the two spindle pole coordinates:

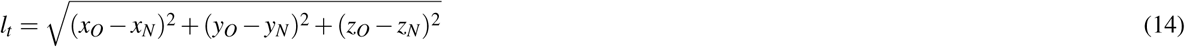

### Kar9 residence time in the bud

The mother-bud boundary is determined by the line defined by the two bud neck coordinates (*x*_1_, *y*_1_) and (*x*_2_, *y*_2_):

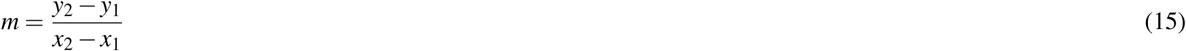

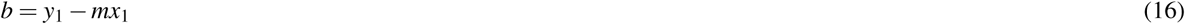

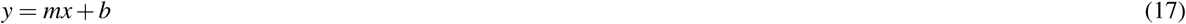

*x,y* coordinates of each Kar9 spot are then used to determine if the coordinates are within the bud region or not. Orientation of the bud is used to establish if the bud is above or below the line defined by the bud neck. For instance, if the polarity axis is between 0*°* and 180*°* (the bud is facing upward), the bud is above the line. The calculation is as follows:

Given the coordinates of Kar9 at a time *t* (*x*_*k*_, *y*_*k*_), the *y*-value corresponding to the *x*_*k*_ on the line of the bud neck is determined:

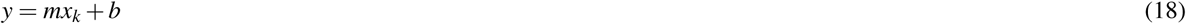

The true Kar9 y coordinate *y*_*k*_ is compared to the calculated y to determine if it is above or below the line of the bud neck: if *y*_*k*_ *≥ y*, (*x*_*k*_, *y*_*k*_) is in the bud, otherwise *y*_*k*_ < *y*, (*x*_*k*_, *y*_*k*_) is in the mother

The number of spots in the bud is recorded and normalized to the length of the acquisition. If the polarity axis is between 180*°*and 360*°* (the bud is facing downward), the inverse comparison relationship is used if *y*_*k*_ *≤ y*, (*x*_*k*_, *y*_*k*_) is in the bud, otherwise *y*_*k*_ *> y*, (*x*_*k*_, *y*_*k*_) is in the mother

### Kar9 depth

Kar9 is projected along the polarity axis as above, where *ϕ* is the polarity axis angle. The midpoint of the bud neck (MPBN) is used as an origin to distinguish Kar9 in the bud (depth *>* 0) from Kar9 in the mother (depth < 0):

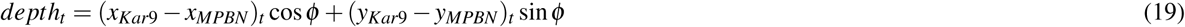

Depth is normalized to the bud size to compare depths between cells:

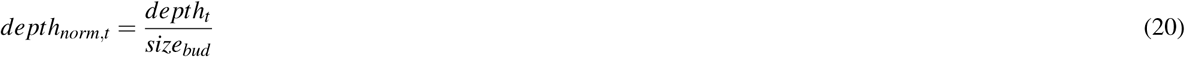

## Acknowledgements

We would like to thank the members of the Vogel lab for fruitful discussions over the course of this project, and Yohann Favire for early contributions to the project. This study was made possible by the Integrated Quantitative Biology Initiative which provided access to the confocal microscope. M. M. was supported by USRA summer fellowship from the Natural Sciences and Engineering Research Council (NSERC). The research was supported by an operating grant from NSERC (RGPIN-2020-05187) and a Canada Foundation for Innovation Innovation Fund awarded to J. V.

## Author contributions statement

M. M. developed the algorithms, performed the experiments and analysis and wrote the manuscript. R. G. Contributed to the development of algorithms and experiments. J. V. conceived of and supervised the project and edited the manuscript.

## Additional information

The authors declare no competing interests.

## Notes

### Competing Interest Statement

The authors have declared no competing interest.

