## supplemental information for "Spindle alignment is uncoupled with Kar9 symmetry breaking"

### Supplementary Materials

#### Supplementary Table S1 – Strains used in this study

| Name | Genotype | Notes | Reference |
|---|---|---|---|
| YV2380 | *Spc42-mKate2-HIS; Kar9-mNeonGreen-NAT; Mat a; LYS(+); met15Δ0(-)* | WT; generated by previous lab member Yohann Faivre. | This study. |
| YV2679 | *Spc42-mKate2-HIS; kar9AA-mNeonGreen-NAT; Mat a; LYS(+); met15Δ0(-)* | *kar9AA* was tagged with mNeonGreen and mated to Spc42-mKate2-HIS by previous lab member Yohann Faivre. | Liakopoulos D, *et al.* (2003); This study |
| YV2412 | *swe1Δ-KAN; Spc42-mKate2-HIS; Kar9-mNeonGreen-NAT; Mat a; LYS(+); met15Δ0(-)* | Made by crossing YV2380 with *swe1Δ* from Invitrogen deletion collection; generated by previous lab member Yohann Faivre. | This study. |

**Supplementary Figure S1 – Spindle elongation is comparable between strains.** Mean spindle growth for WT (grey), *kar9AA* (blue) and *swe1Δ* (pink). Distributions are comparable between strains. Spindle growth is independent of spindle length (mean and initial lengths).

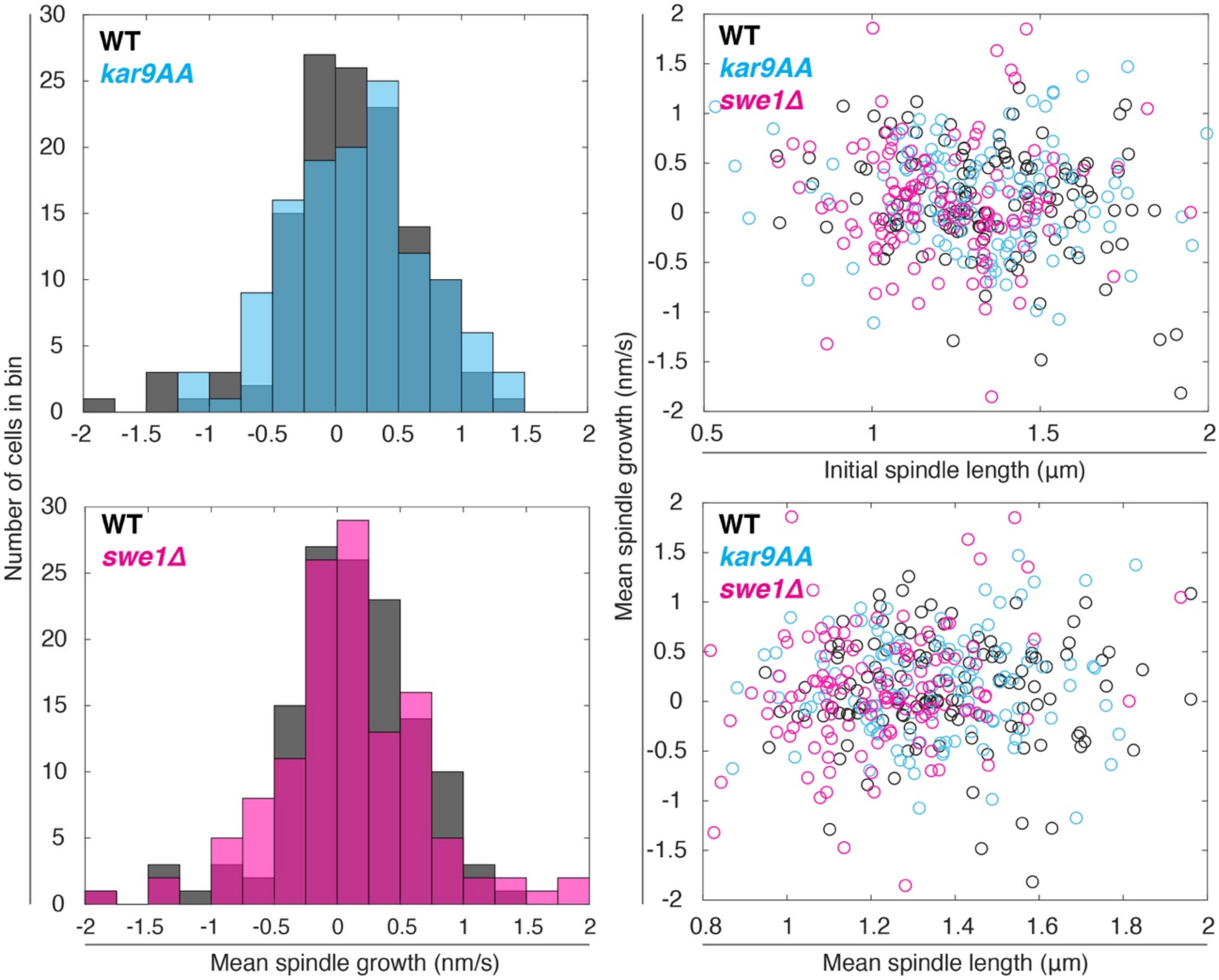

**Supplementary Figure S2 – 2D alignment measures.** (A-C) Symmetric spindles are unable to align themselves, regardless of genotype. (D) 2D alignment of WT spindles is biased toward small angles (proper alignment). (E) A subset of asymmetric *kar9AA* spindles are moderately biased toward small angles, while the others are clustered about misalignment (90$^o$). (F) Asymmetric *swe1Δ* spindles are biased toward short angles but less efficient than WT asymmetric spindles. All are consistent with 3D measures described in the main text.

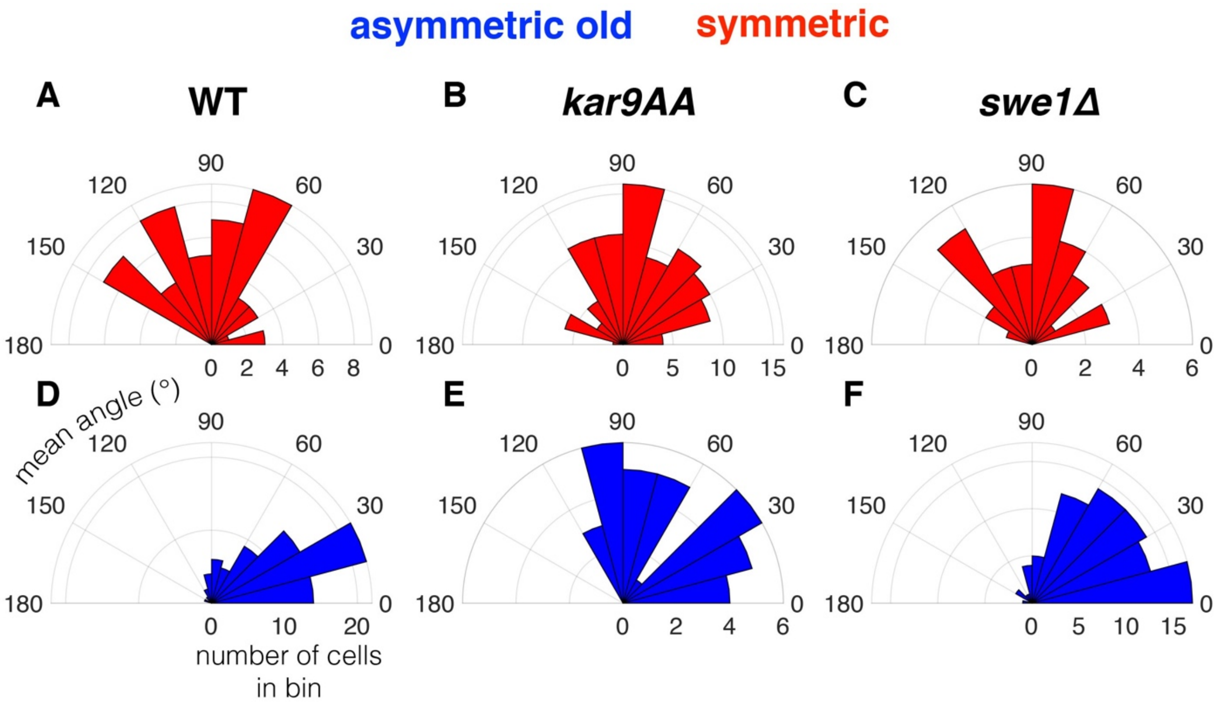

**Supplementary Figure S3 – Kar9 depth is correlated with spindle alignment.** (a, inset) Graphical description of Kar9 depth measure. (a-c) Kar9 depth of PA and NA spindles of WT(a), *kar9AA*(b) and *swe1Δ*(c) cells. Regardless of genotype, Kar9 of PA spindles preferentially reside in the bud and surrounding the bud neck. Kar9 of NA spindles explore more space, ranging from the distal-most part of the cell to within the bud. (d-f) Alignment index is proportional to Kar9 depth and Kar9 residence time in the bud, irrespective of symmetry state.

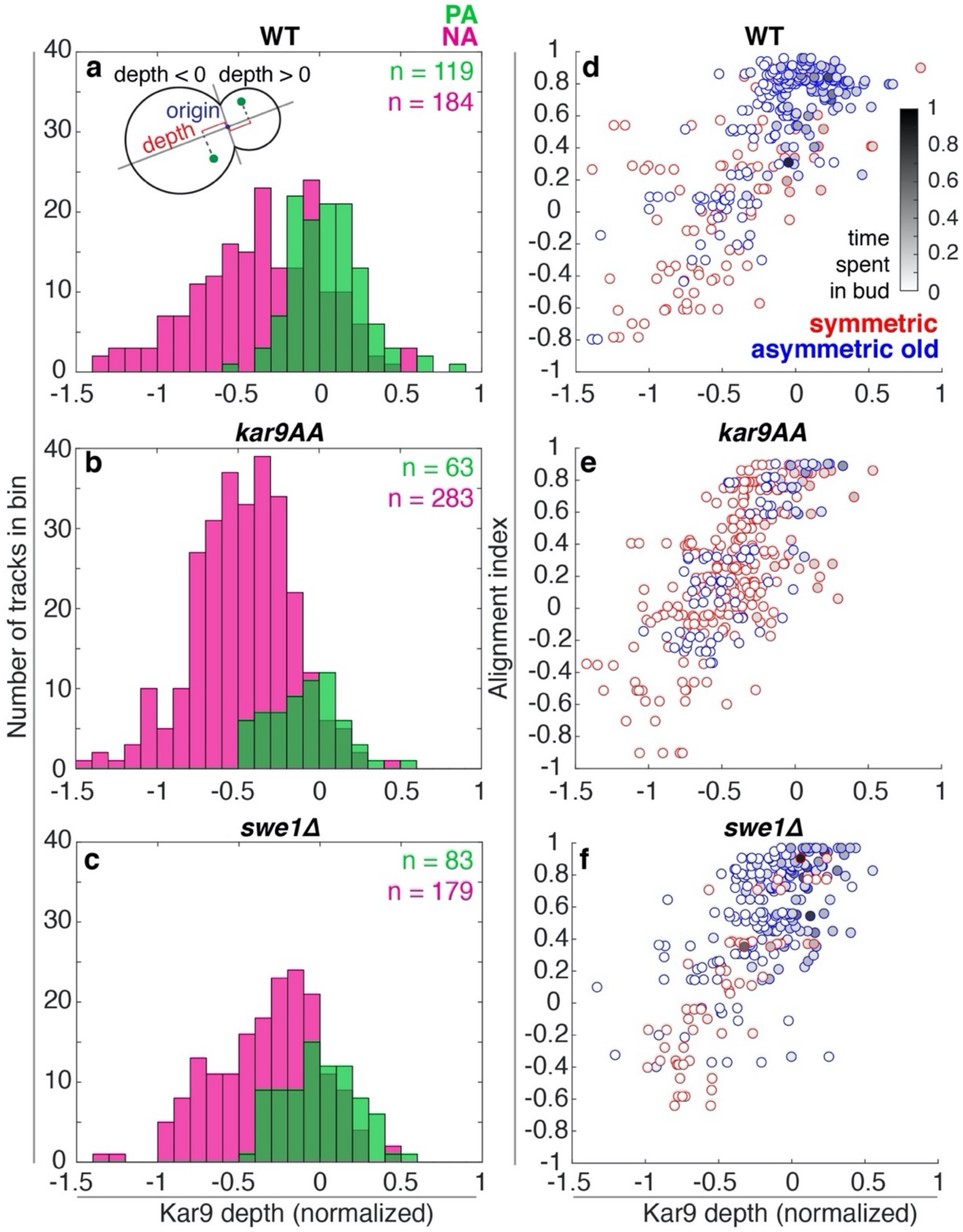

**Supplementary Figure S4 – Cumulative fraction of cells equal to or above a given Kar9 residence time threshold.** (a-c) Same data as show in Figure 5d-f, with the addition of classes of cells with sample sizes below 5 cells.

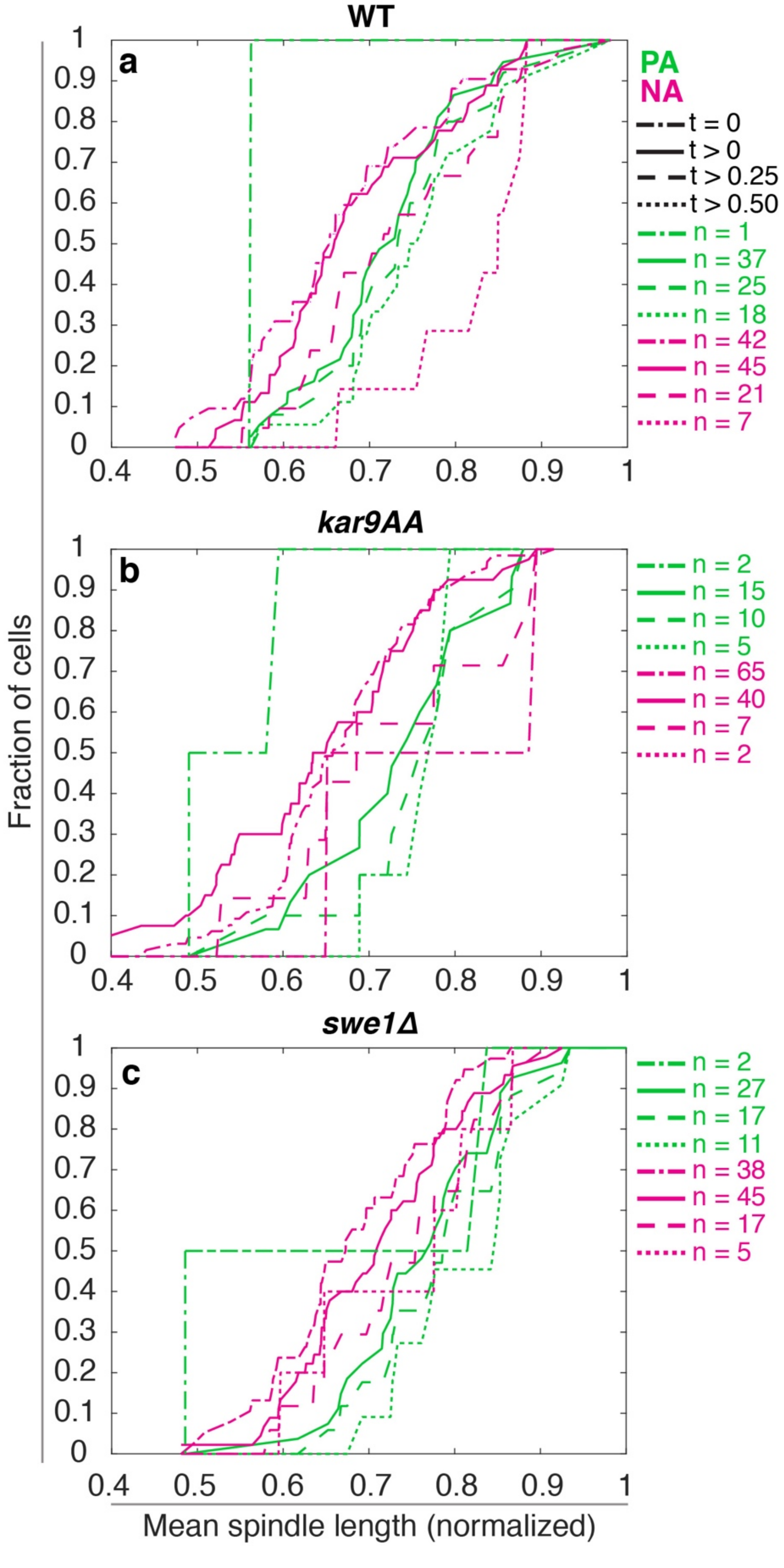
